# The interoceptive hippocampus: mouse brain endocrine receptor expression highlights a dentate gyrus (DG)–cornu ammonis (CA) challenge– sufficiency axis

**DOI:** 10.1101/802975

**Authors:** Richard Lathe, Sheena Singadia, Crispin Jordan, Gernot Riedel

## Abstract

The primeval function of the mammalian hippocampus (HPC) remains uncertain. Implicated in learning and memory, spatial navigation, and neuropsychological disorders, evolutionary theory suggests that the HPC evolved from a primeval chemosensory epithelium. Internal sensing deficits in patients with HPC lesions argue that internal sensing may be conserved in higher vertebrates. We studied the expression of 250 endocrine receptors in mouse brain. Key findings are (i) the proportions and levels of endocrine receptor expression in the HPC are significantly higher than in all other comparable brain regions. (ii) Surprisingly, the distribution of endocrine receptor expression within mouse HPC was found to be highly structured: receptors signaling ‘challenge’ are segregated in dentate gyrus (DG), whereas those signaling ‘sufficiency’ are principally found in *cornu ammonis* (CA) regions. Selective expression of endocrine receptors in the HPC argues that internal sensing remains a core feature of hippocampal function. Further, we report that ligands of DG receptors predominantly inhibit both synaptic potentiation and neurogenesis, whereas CA receptor ligands conversely promote both synaptic potentiation and neurogenesis. These findings suggest that the hippocampus acts as an integrator of body status, extending its role in context-dependent memory encoding from ‘where’ and ‘when’ to ‘how I feel’. Implications for anxiety and depression are discussed.

## Introduction

Current thinking predominantly attributes to the hippocampus (HPC) a pivotal role in learning and memory, in spatial navigation, and in anxiety, stress, and depression. However, the central function of the HPC in both memory and neuropsychological disorders may be consistent with an underlying role in internal sensing (interoception). Previous studies have implicated cortical regions, limbic brain, and thalamus, as well as the hypothalamus and brainstem regions, among others, in interoception [1]. The HPC (and adjoining amygdala) is a prominent contender – in addition to his profound learning and memory deficits following HPC surgery to alleviate severe recurrent epilepsy [2], the famous patient H.M. was unable to sense internal states [3]. Similar observations have been made in rodents with selective HPC lesions [4–6].

A role for the HPC in internal sensing is consistent with evolutionary theory that the HPC (and olfactory system) arose from a chemosensory epithelium, but with the closing of the brain ventricles during evolution the hippocampus retained the capacity to sense the internal milieu of the body [7–9]. It is of note that the ‘rostral migratory stream’ in neonatal mice directly connects the HPC and the chemosensing olfactory system [10], consistent with a common developmental origin. In addition, a key characteristic of traditional sensory epithelia such as the olfactory system and retina in many vertebrate species is that neurogenesis continues into adulthood [11,12], and neurogenesis is also prominent in adult hippocampus, principally underlying the dentate gyrus (DG) (reviewed in [13].

Internal sensing is a key modulator of behavior. Hunger and thirst are induced by deficiencies in nutrient and water, respectively, and elicit clear adaptive motivations and behaviors. Other diverse internal states, ranging from salt deficiency to hormonal status to inflammation/infection, exert powerful effects on multiple aspects of brain function, centrally including adaptive behavior as well as learning and memory, but the target brain region(s) and receptors remain poorly defined.

The anatomy of the mammalian HPC is consistent with an internal sensory role. The hippocampal formation lies at the interface (*limbus*, ‘fringe’) between the lower brain and the mass of the cerebral cortex. In terms of blood supply, the HPC is perhaps the most highly irrigated of all brain regions, and is also flanked by the central and lateral ventricles with the choroid plexus [14]. In cross-section, the formation is divided into CA regions CA1 and CA3 (with a short intervening structure, CA2), and the DG. There may be a further functionally distinct region, the dentate hilus, but this is less secure. Gene expression surveys largely confirm this anatomy [15,16]. Some have introduced additional subdivisions both within the DG–CA circuit [17] and along the length of the hippocampus [18]. However, for simplicity we retain the conventional subdivisions CA1–CA3 and DG.

To address the physiological role of the HPC we previously employed differential hybridization [19], candidate gene screening [20], and gene-trapping [21] to identify genes selectively expressed in HPC. This revealed that the mouse HPC expresses several receptors and signaling molecules, potentially indicating a role of the HPC in internal sensing of body physiology [9]. The aim of the present study was therefore to test rigorously the hypothesis that the hippocampus is involved in interoception through systematic analysis of the expression patterns of endocrine receptors across mouse brain, including subregions of the HPC.

Specifically, we sought to answer two central questions. (i) Does the mouse HPC express a greater diversity and/or level of endocrine receptors than other brain regions such as the cortex and the cerebellum? (ii) If a greater level of expression is found, are these receptors expressed uniformly across the HPC, or are different receptors differently distributed in the different subdivisions of the HPC? – and can the pattern of expression tell us anything about the function of the HPC? We report that the HPC is the principal brain site of endocrine receptor expression and, perhaps surprisingly, this analysis revealed a highly segregated distribution of receptor expression in mouse hippocampus.

## METHODS

### Endocrine receptors

A list was assembled of receptor molecules in mice and humans that respond to endocrine (blood-borne) ligands. We elected to study 250 receptors, a number chosen to minimize the risk that a small number of atypical receptors or experimental artifacts might bias the overall picture, weighed against the labor-intensive constraints of manually analyzing a larger number of receptors. To assemble the list, the GeneCards database (www.genecards.org) was searched at random for genes/gene products containing ‘receptor’. A preliminary list (>>250 receptor candidates) was manually filtered to exclude (i) non-receptor entries (e.g., receptor downstream kinase, etc.), (ii) evident receptors for neurotransmitter and non-diffusible cell–cell interaction molecules, and (iii) receptors not listed in the primary database consulted (Allan Brain Atlas) as well as receptors whose expression profiles were classified as failing quality control. Although principally cell-surface molecules, the final list includes intracellular receptors with an endocrine role (e.g., nuclear receptors). This generated a list of 253 endocrine receptors (Table S1 in the supplementary material online; the molecular functions of specific groups of receptors are discussed in Box S1).

### Quantification of mouse brain endocrine receptor expression data

Primary analysis relied on the Allen Brain Atlas (ABA; http://mouse.brain-map.org/), a publicly available repository of *in situ* hybridization gene expression data across mouse brain [22] made available by the Allen Institute for Brain Science established by Paul G. Allen. To retrieve expression patterns we entered search terms (e.g., Gene1) into http://mouse.brain-map.org/search/show, sagittal sections were selected in all cases when these were available. The ‘expression’ option and the target brain region (typically mid-brain including the hippocampus) were selected, a screenshot was taken; data for all 253 receptors were recorded at the same magnification and intensity in a repository of image files. To quantitate expression levels ImageJ [23,24] was employed. Using default settings, and a standard image size, representative brain regions (HPC; cortex, CX; and cerebellum, CB) were selected using a cursor box of constant size and analyzed using the ‘measure’ function of ImageJ (the olfactory bulb could not be systematically analyzed because this structure can be lost during dissection, and the small relative size of the mouse hypothalamus precludes analysis at the resolution afforded by ABA). In each case the ‘Mean’ function was used instead of the integrated density function ‘IntDen’ because, at constant image size, the relative values are the same. The same technique was used for hippocampal subregions, but the cursor box was manually fitted to the separate regions (CA1, CA2, CA3, DG). The ‘Mean’ function in these cases represents relative (total) expression of the target gene within the region measured. These analyses generate a digital intensity reading on a scale of 0 to 255. The program accommodates different colors as follows: black, 0.00; red, 85/255 (0.333); yellow, 170/255 (0.666); white, 255/255 (1.000), mirroring the output of the ABA. Because region selection is to some extent subjective, subregion expression analysis was performed by two independent researchers; in cases of disparity consensus was reached following reanalysis of the primary data. Values were then normalized – a biologically realistic data transformation because (i) the signal for each target depends on the hybridization properties of the specific probe employed, (ii) the biological effects of a given receptor will vary across a wide range depending on ligand concentration, ligand affinity, and downstream signal transduction, and (iii) for a given gene, the inter-regional pattern (ratio) of expression across the brain/hippocampus (unlike absolute values) is likely to be independent of the specific probe/hybridization parameters. For normalization, the highest expression value was selected (100%) and expression in other regions was expressed as a percentage of maximum. Inter-region expression ratios in whole brain were calculated from the un-normalized expression data. Primary data for receptor gene expression across the brain are given in supplementary Table S2, and for hippocampal subregions in supplementary Table S3.

### Heat mapping and statistical analysis

All analyses focused on genes that were expressed in at least one of the selected brain regions (98 genes in Figure 1, and 86 genes in Figure 2), and were conducted in the R programming environment, version 3.3.3 [25]. Heatmaps were generated using heatmap.2 in the gplots library for R. Note that heatmap.2 provides dendrograms to aid visualization of relationships among components of the heatmap but provides no statistics to indicate support for the presented dendrograms versus alternative, competing dendrograms. Therefore, we strongly caution against overinterpretation of the dendrograms presented.

**Fig. 1.**
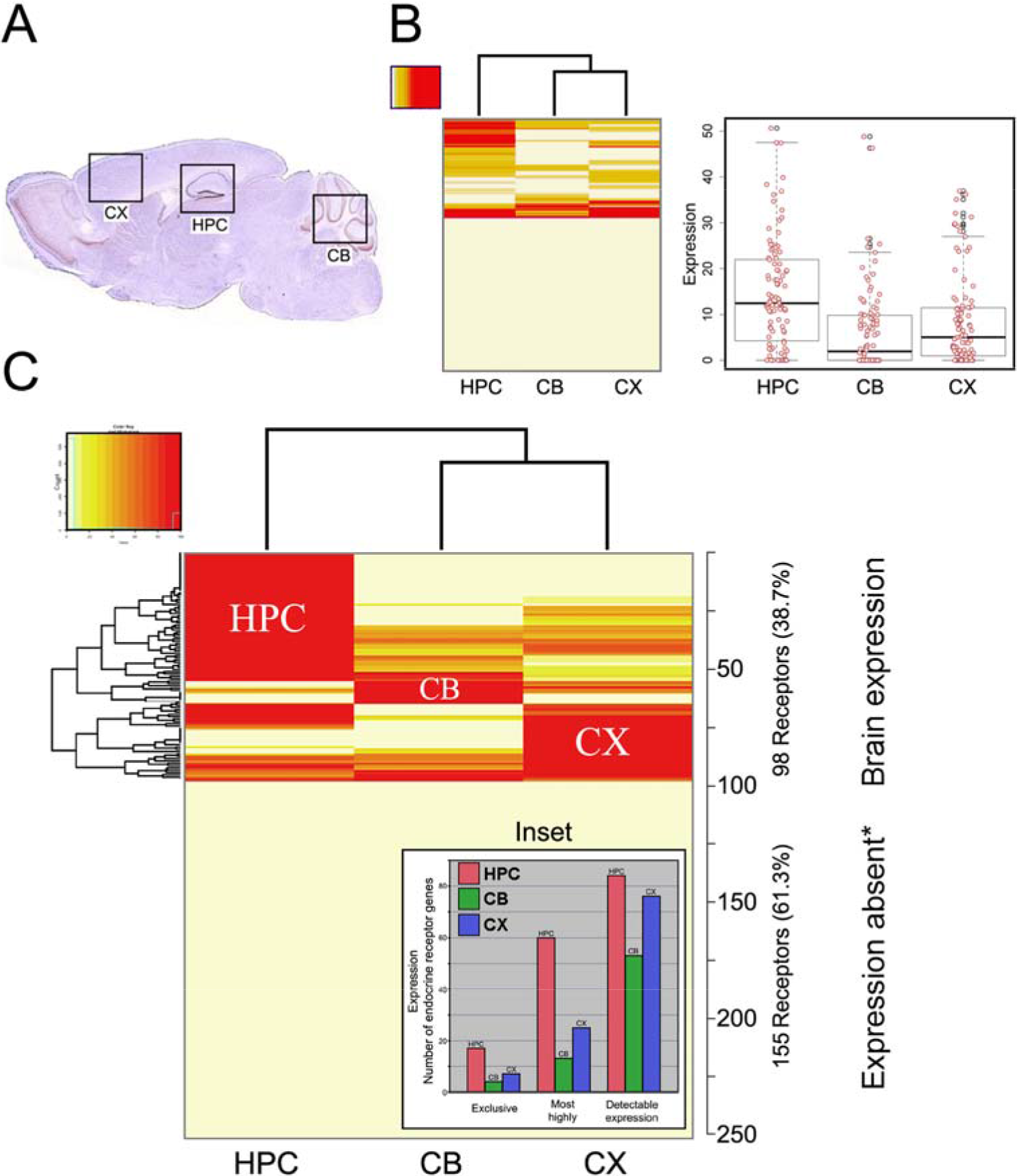
Endocrine receptor gene expression in mouse brain and enrichment in the hippocampus (HPC). More than one third of all endocrine receptors were detectably expressed in brain, where they are likely to modulate brain function and cognition. Expression was restricted to specific brain regions: other than hippocampus (HPC), cerebellum (CB), and cortex (CX), there was little evidence for specific gene expression in other comparable regions (~4%). (**A**) Mouse brain section highlighting the three regions studied in detail: HPC, CB, and CX. (**B**) (Left) Heatmap of ‘raw’ (unnormalized expression data, see Methods) for HPC versus CB and CX. (Right) Scatterplots of unnormalized expression levels; horizontal lines are medians and quartiles showing that the mean expression level of all receptors in HPC was significantly higher than in either CB or CX. (**C**) Normalized (maximum expression level = 100%) gene expression data. On three counts, the HPC (red), versus CB (green) and CX (blue), is the major site of expression of endocrine receptors (253 receptors examined) as further evidenced by the inset showing (i) exclusive expression in HPC, (ii) most prominent expression in HPC, (iii) overall number of receptors expressed. *Receptors showing no detectable expression or low-level/punctate/irreproducible expression are classified as expression absent. Note that the dendrograms (generated by heatmap.2), depicted in A and B, are not supported by statistical analysis versus alternative, competing dendrograms. Genes that are expressed exclusively in HPC, CB, or CB were not distributed among these three regions with equal probability, and ‘exclusive genes’ were expressed most often in HPC; the same result emerges when considering genes that are expressed most prominently in one brain region. Thus, the HPC expresses both a greater number and level of endocrine receptor genes than any other brain region analyzed.

**Fig. 2.**
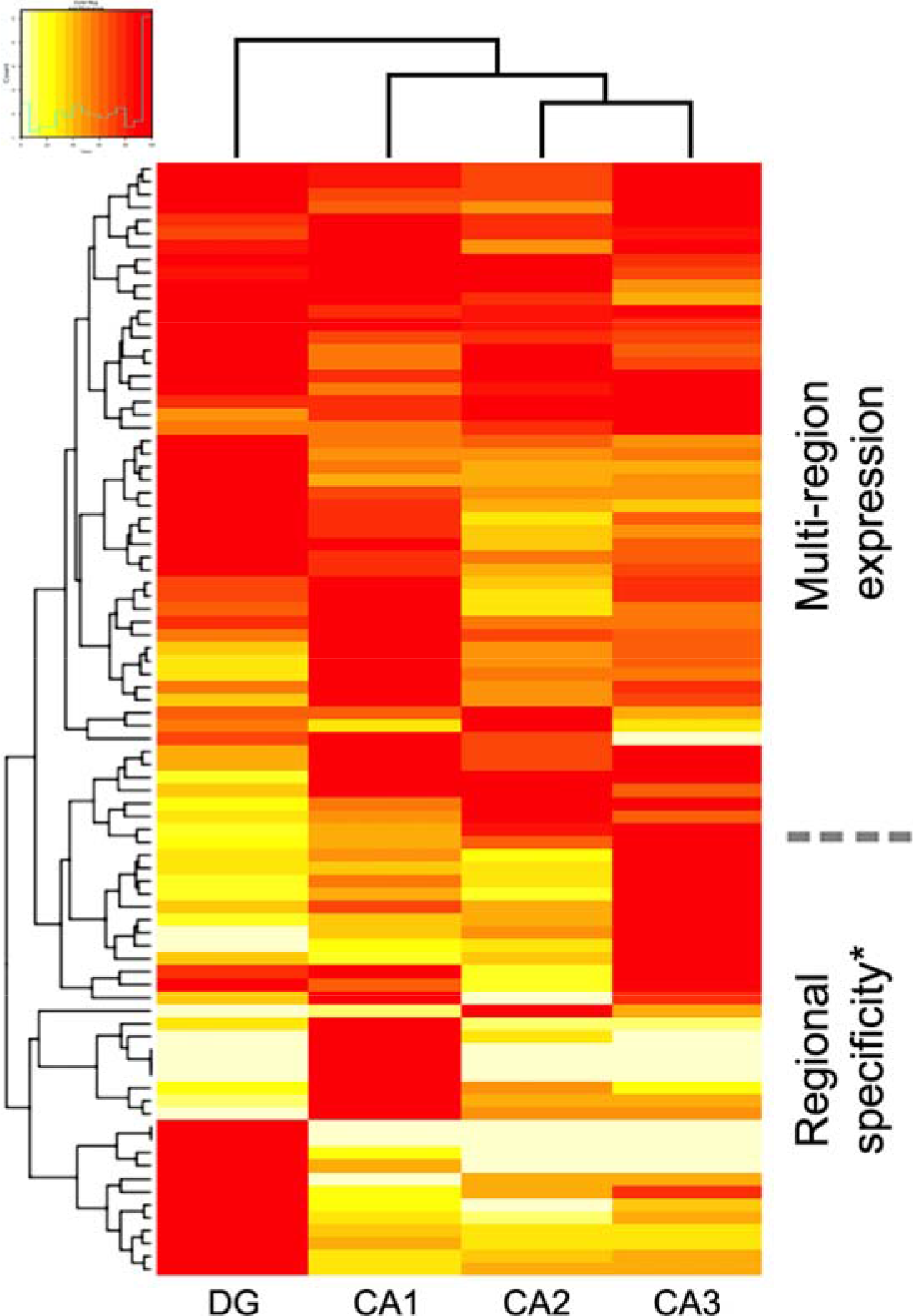
Subregional representation of 86 receptors expressed in mouse hippocampus (HPC). Data are normalized to the maximum expression level. *The data indicate that some receptors are somewhat restricted in their expression pattern to one subregion, whereas others are expressed in combinations of regions. Note: the depicted dendrograms (generated by heatmap.2) are not supported by statistical analysis versus alternative, competing dendrograms. There were significant positive correlations between CA2 and CA3, and significant negative correlations between CA1 and DG (Table S4).

To test whether gene expression profiles differ across brain regions (HPC, CX, and CB) we measured the correlation in gene expression among brain regions. To this end, we analyzed normalized gene expression (see above) because variation in probe affinity may generate spurious correlations. We calculated the correlation using arcsine square root transformed values of normalized gene expression, and used case-bootstrapping to generate 95% confidence intervals (R package ‘boot’ [26,27]; bootstrapped 10 000 replicates).

Wilcoxon signed rank tests and paired *t* tests were used to determine whether non-normalized gene expression differed among brain regions (Wilcoxon tests to compare HPC, CX and CB; paired *t* tests to compare CA1, CA2, CA3, and DG). We used Chi-square goodness of fit tests to determine whether genes that are exclusively (or alternatively, predominantly) expressed in HPC, CB, or CX are distributed equally among these regions. We used a series of three binomial tests to determine whether the numbers of genes expressed differed among HPC, CB and CX. Pairwise correlation analysis is given in Table S4.

### Informative genes

For the majority of receptor genes the biological function of the receptor and/or the identity of the ligand(s) remains unknown. For further analysis we therefore selected an ‘informative’ subset of 32 genes where information is available concerning the biological role (or inferred role) of the ligand/receptor pair. This subset included receptors for known diffusible hormones (e.g., estrogen, glucocorticoids, progesterone), for cytokines (e.g., interleukins, interferons, tumor necrosis factor), and growth factors (e.g., fibroblast growth factor). The list of informative genes is presented in Table S5.

### Inter-region expression ratios in hippocampus; statistical analysis

Normalized expression data were used to test whether gene expression ratios among hippocampus regions differed between group A versus B genes (for an explanation of groups A and B see Results and Discussion). The mean expression data for CA (CA1–3) and DG were calculated and then log-transformed (1 or 2 was added to all values prior to log-transformation to account for zeros; the outcome was the same in both cases). Pairwise DG/CA expression ratios (Δ) were calculated from Δ = log(DG) − log(CA) {therefore, Δ = log(DG/CA)}. Welch’s *t* test was employed to assess statistical significance of pairwise differences in ratios (i.e., Δ) for informative (group A, challenge; and group B, sufficiency) genes. The same approach was employed for HippoSeq data (below).

However, because the distribution of Δ violates the assumptions of *t*-tests, we additionally used a permutation test to confirm conclusions from the *t*-test. The permutation test has two stages. First, average Δ was calculated for each group of A and B genes, and the difference between these averages was calculated. This value represents the observed difference in average Δ between groups A and B. Second, (i) Δ values were randomized among groups A and B, (ii) average Δ of these randomized data was calculated for group A and B genes, and (iii) the difference between these average Δ values between groups A and B was calculated. We repeated this second stage 10 000 times to generate a null distribution, against which we compared the observed difference in average Δ between group A and B genes to yield the *P* value reported here.

We used Dunn–Sidak corrected critical *P* values to assess significance when making multiple comparisons (*P*_crit_ = 0.0169 and 0.00851 for three and six comparisons, respectively).

### Cross-validation of expression data

To validate data from the Allan Brain Atlas we consulted HippoSeq (https://hipposeq.janelia.org) [28], a database of gene expression data. HippoSeq is based on transgenic tagging of subregions of mouse HPC, brain microdissection, fluorescence cell-sorting retrieval of target HPC CA pyramidal cell/dentate neuronal populations, and deep sequencing of mRNA populations. A revised input format (kind courtesy of Cembrowski *et al.*) allowed query of multiple genes, generating a table of absolute readcounts (FPKM, fragments per kb of transcript per million mapped reads). Cross-comparison to ABA established a lower limit (null expression) where 4 FKPM equated to an undetectable hybridization signal (not presented). Parallels and differences between the ABA and HippoSeq studies are summarized in Table S6.

Because ABA is more robust than HippoSeq in terms of the number of animals studied (a small number of unrepresentative animals would be less likely to affect conclusions based on ABA rather than on HippoSeq, Table S6), and because an *in situ* hybridization pattern (ABA, particularly if confirmed by identical patterns generated in other mouse strains or species) may be more immune to bias than an automatically generated value (HippoSeq), ABA was preferred over HippoSeq for our primary analysis, although both are reported where appropriate.

### Analysis of receptor function

PubMed was searched for the name of each individual receptor in conjunction with ‘synaptic potentiation’ OR ‘synaptic plasticity’ OR ‘long-term potentiation’ OR ‘LTP’ OR ‘neurogenesis’. Relevant publications were manually tabulated for ligand effects on both parameters and are listed in Table S8. Intergroup pairwise comparisons of effects (inhibition versus stimulation) of literature-recorded ligands on LTP and neurogenesis employed both Student’s unpaired *t* test and chi-square test.

## RESULTS

A representative list of 253 endocrine receptors was compiled (neurotransmitter receptors and cell–cell interaction molecules were excluded; supplementary Table S1). *In situ* hybridization patterns were extracted from the Allen Mouse Brain Atlas (ABA); these were manually scanned and quantified (Methods). Where appropriate, values were normalized and inter-region ratios calculated.

### Brain distribution of endocrine receptor expression

We report that, of all endocrine receptors, 98/253 (38.7%) were detectably expressed in brain. This argues that, in addition to regulating body physiology including growth, development, reproduction, and homeostasis, etc., a major proportion of endocrine receptors may directly regulate brain function and cognition.

We also report that endocrine receptor expression in mouse brain is limited to specific brain regions. Only a small number of genes were expressed in major areas such as the olfactory bulb (OLF), thalamus, pons/medulla, pallidum, or striatum (4.3%; see below). This focused our attention on HPC, cortex (CX), and cerebellum (CB). Hypothalamus could not be examined (Methods and Discussion).

Regarding our first question – the proportion of endocrine receptors expressed in mouse HPC – we report that 86 of 253 (34.0%) endocrine receptors are expressed in HPC, a higher number than in either CB (53) or CX (76). Importantly, the level of expression in was highest in HPC. Of all receptors with detectable expression in brain (*n* = 98), 61.3% were most prominently expressed in the principal neuronal layers (pyramidal and granule cells) of the HPC (versus 9.1% in CB and 25.5% in CX). Indeed, 17.3% of brain-expressed endocrine receptors were exclusively expressed in HPC (compared to 4.1% and 7.1% that were exclusively expressed in CB and CX, respectively). Fig. 1 presents heatmaps of the normalized and un-normalized expression data for these three brain regions, and the inset gives numerical values for exclusivity, most prominent, and detectable expression.

Non-normalized gene expression differed significantly in all pairwise comparisons among HPC, CB, and CX. The HPC expressed these genes at significantly higher levels than either CX or CB (Wilcoxon signed rank test; vs CX: *V* = 3467, *P* = 3.266e−06; vs CB: *V* = 3527, *P* = 1.398e−08), and CX expressed genes at higher levels then CB (*V* = 2208.5, *P* = 0.009944). All comparisons remained significant after accounting for multiple comparisons. Overall, the probability of detectable gene expression was significantly higher for HPC than CB (binomial test, *P* = 0.0101), but did not differ for remaining comparisons (binomial tests; HPC and CX: *P* = 0.58; CX and CB: *P* = 0.052); these results remain unchanged after accounting for multiple comparisons.

Although we were unable to systematically screen for expression in OLF (Methods), a very small number of genes from our selection were expressed in OLF (*Ednrb*, *Epor*, *Ccr3*, *Crhr1*, *Nrp1*, and *Nmbr*) of which only *Ccr3* and *Nmbr* appeared to be specific for OLF. Remaining genes were expressed in striatum and/or pallidum (*Acvrl1*, *Nfgr*, *Rarb*) or in pons/medulla (*Adipor2*, *Esrrg*). No other brain regions stood out with other than trace expression in this survey (small foci of low-level expression, not presented); in total, these represent 4.3% of all the endocrine receptors studied, a far lower proportion than in either HPC, CB, or CX.

We conclude that, based on 253 receptors, there is significantly greater endocrine receptor gene expression in HPC than in either CB or CX, or in any other comparable brain region analyzed (noting that hypothalamus could not be studied; Discussion).

### Distribution across hippocampal subregions

With regard to our second question – the pattern of expression within the HPC – all the receptors studied with detectable HPC expression (*n* = 86; Fig. 1) identified mRNA within the cell bodies of the principal excitatory neurons (pyramidal cells, DG neurons) of the HPC. However, the expression patterns of the assembled genes were non-randomly distributed across subregions – although some were detectably expressed in all subregions, many were expressed only in restricted regions of the HPC. Fig. 2 presents the distribution (heatmap) of receptor expression across the different regions of the mouse HPC. To address correlations between HPC subregions, we performed pairwise correlation analysis (Table S4). Normalized gene expression was significantly negatively correlated between DG and CA1, and positively correlated between CA2 and CA3. All remaining combinations of CA1, CA2, CA3, and DG provided no evidence of correlated gene expression (Table S4).

To validate the subregional distributions in mouse HPC, we compared ABA *in situ* hybridization data against a second database, HippoSeq (Methods; this database only addresses HPC expression). The HippoSeq database supported the overall subregional expression patterns detected by *in situ* hybridization.

### Distribution of receptors with established roles: subregion–function correlations reveal a challenge–sufficiency axis

For the majority of the receptors studied here the biological ‘meaning’ is unknown, either because the receptor ligand is unknown or because the physiological role of the ligand(s) has not been established. To illustrate, the first and last genes in our list, *Acvr1* and *Vmnr234*, respectively encode activin A receptor type 1 and a vomeronasal-like receptor. Ligands for ACVR1 include both inhibins and activins, that inhibit and activate diverse physiological processes and, moreover, have opposing functions; the primary *in vivo* ligand for ACVR1 in the CNS remains unknown. For VMNR234, the ligand is also unknown. Given this uncertainty we examined receptors from an ‘informative’ list (*n* = 32) where the function of the ligand is known (or inferred): these include angiotensins, cytokines, fibroblast growth factor (FGF), interleukins/interferons, prostaglandins, retinoids, steroid hormones (androgens, estrogens, glucocorticoids and mineralocorticoids), tumor growth factor (TGF), and tumor necrosis factor (TNF) (Methods and Table 3). This revealed a gradient of expression across the HPC, where some receptors were principally expressed in DG regions, and others were principally expressed in CA regions (Fig. 3).

**Fig. 3.**
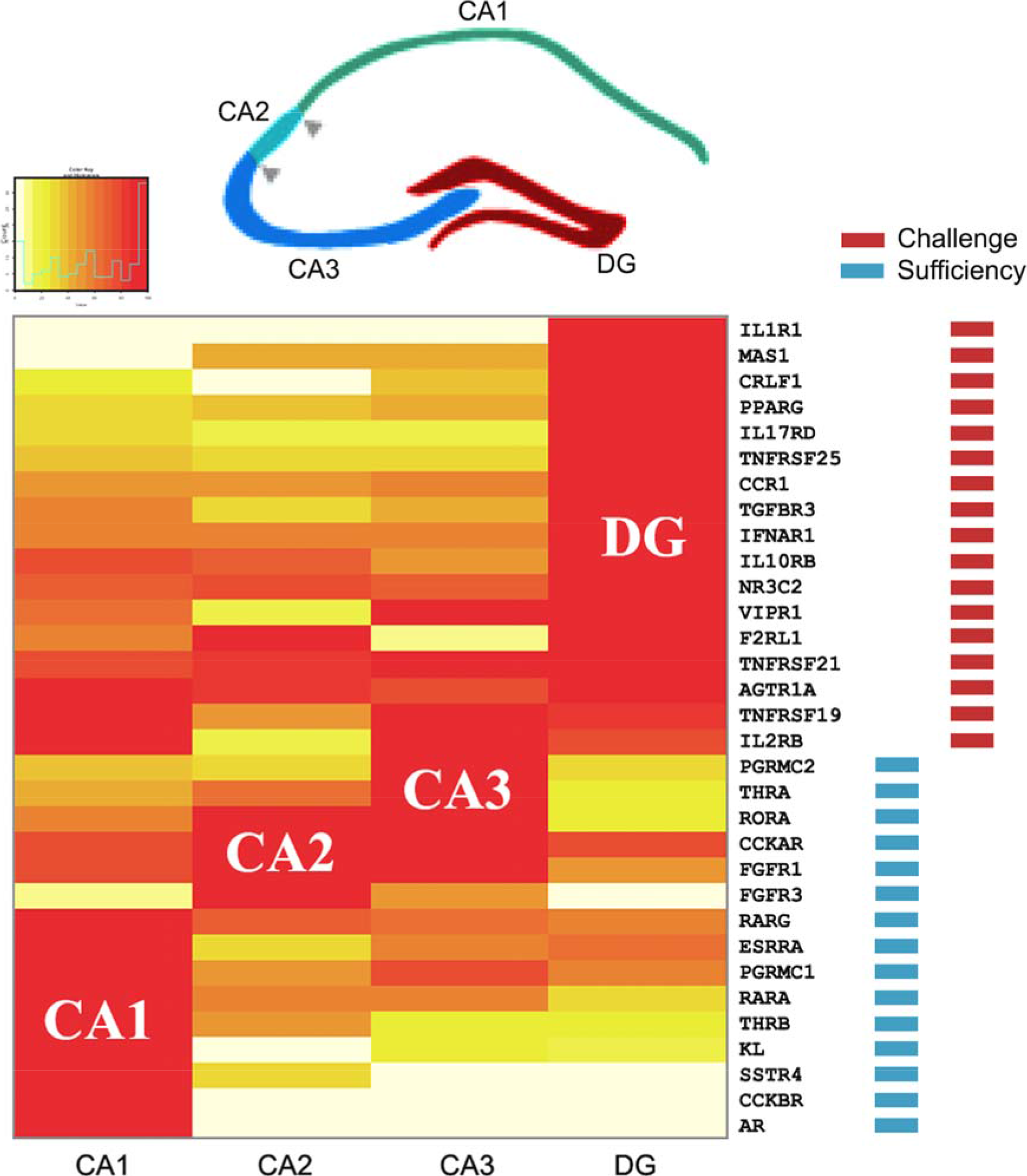
Expression of ‘informative’ endocrine receptors in subregions of the mouse hippocampus (HPC). (Above) Principal neuroatomical subdivisions of the rodent HPC (adapted from the model of [15]). (Below) Informative (see main text) receptors sorted according to regional expression (heatmap, normalized data) with CA1 and DG at the two extremes (Methods) showing expression clustering of receptor types in different regions (e.g., ‘sufficiency’ – FGF receptors FGFR1, FGFR3, and KL in CA regions; and ‘challenge’ – interleukin and TNF receptors IL1R1, IL17RD, IL10RB, IL2RB, TNFRSRF 25, TNFRSF21, TNFRSF19 in DG).

### Receptor categorization by function

We sought a unifying principle that might underpin and explain the gradient of receptor expression. It became apparent that receptor function differed according to location within the HPC. Receptors reflecting stress of various types (e.g., receptors for inflammatory cytokines and glucocorticoids) provided a clue because their expression was clustered in DG.

Conversely, it was noted that receptors responding to growth-promoting ligands (e.g., growth factors and sex steroids) were principally localized in CA regions. On this basis it was possible to classify each ligand/receptor pair into two groups.

Because one group of receptor ligands (designated ‘group A’) signal loss of homeostasis and/or physiological stress of various types (these ligands include angiotensins – blood pressure fall; glucocorticoids – stress hormones; cytokines, interferons, and TNF – immune challenge), we describe these here as denoting ‘challenge’, whereas a second group of ligands (‘group B’) conversely includes growth-promoting hormones and factors (e.g., androgens, estrogens, fibroblast growth factor, retinoids), which we term here ‘sufficiency’ (more detailed listing and discussion of receptor function is presented in Table S8 and Box S1). Although this classification is fully open to debate and refinement, we believe that it provides a potential interpretation of the observed gradient of expression.

As shown in Fig. 3, there was unexpected clustering of group A (‘challenge’) receptor expression in DG, whereas group B (‘sufficiency’) receptors were predominantly expressed in CA regions.

To address the statistical significance of the patterning of group A versus group B observation we calculated the ratios between different hippocampal subregions (mean of CA regions versus DG) by conversion to log_10_ values and subtraction (Methods) and plotted the results for the two groups A and B (Fig. 4). The ratio of gene expression in CA to DG differed significantly between group A and B genes (Welch’s *t*-test, *t* = 4.22, *df* = 27.69, *P* = 0.00024). The same analysis was then repeated for the HippoSeq data; this also achieved significance for CA regions versus DG (*P* = 0.0016). Permutation tests confirmed these findings.

**Fig. 4.**
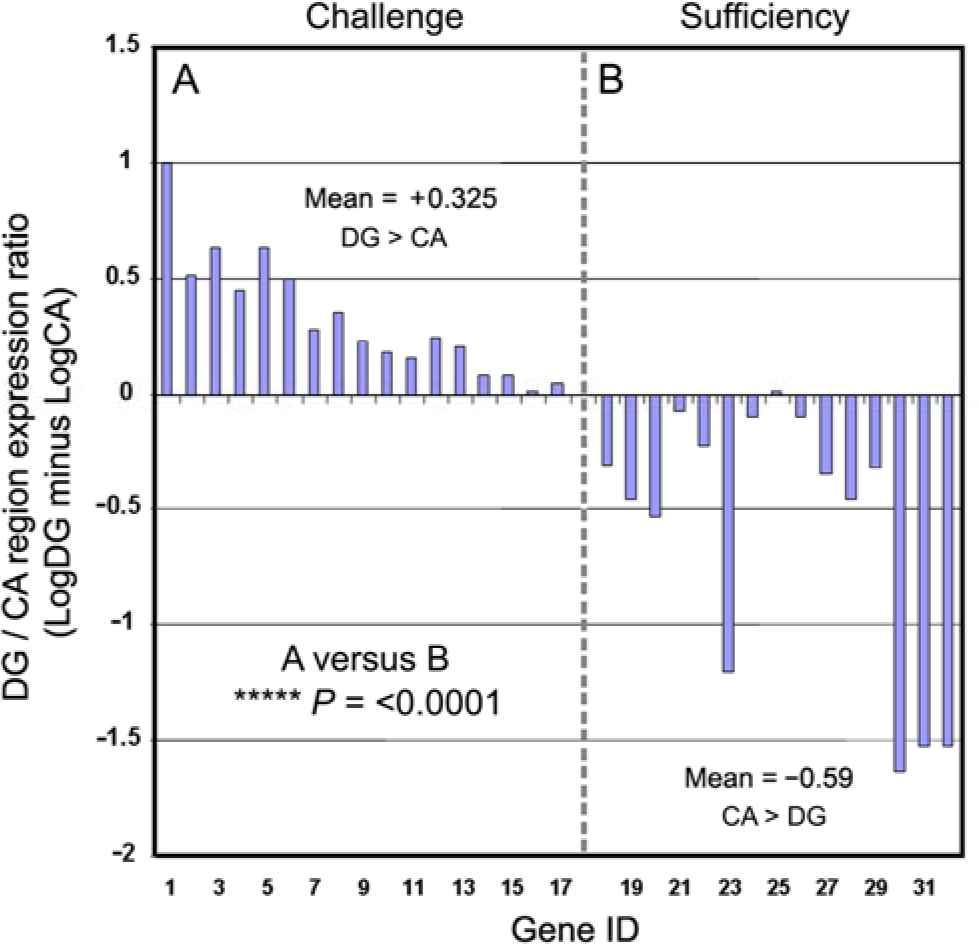
Ratios of CA versus DG expression for informative receptors. (**A**) Group A (DG/challenge). (**B**) Group B (CA/sufficiency). Individual genes are ordered as in Fig. 3. The differential DG versus CA pattern of expression was highly significant.

We conclude that the expression pattern is highly structured within mouse HPC, and that group A receptors (‘challenge’) are preferentially expressed in DG, and group B receptors (‘sufficiency’) are selectively expressed in CA regions (Fig. 3).

### Further receptors confirm the generality of the axis

To test whether the axis extends to other endocrine receptors, we examined the expression pattern (in both ABA and HippoSeq) of other informative receptors (that were not on our original list) whose ligand is known and that are expressed in brain. We identified five such receptors. All were expressed in mouse HPC (although interleukin 6 receptor was only expressed at low levels, Table S7). Challenge receptors (interleukin 6 receptor, growth hormone secretagogue receptor, opioid growth factor receptor, and irisin receptor) were all expressed at higher levels in DG than in CA regions, whereas sufficiency receptors were expressed at highest level in CA regions (glucagon-like peptide 1 receptor) or were expressed at similar levels in CA and DG (leptin receptor) (Table S7), confirming (5/5) that the DG versus CA differential ratio extends to other receptors, reinforcing the generality of our findings.

### HPC receptors are functional: synaptic potentiation and neurogenesis

We addressed whether the informative receptors are functional *in vivo* and *in vitro* by literature searching regarding two output measures: synaptic potentiation (long-term potentiation, LTP) and neurogenesis. The evidence argues that these endocrine receptors are fully functional and modulate both LTP and neurogenesis.

#### Synaptic potentiation

Although not all receptors have been studied in the literature, there was evidence that DG ligands predominantly inhibit LTP, whereas CA ligands promote LTP. For example, DG ligands IL-1, IL-2, IFN-α, IFN-γ, TGF-β, and TNF-α all inhibit LTP in rodent hippocampus [29–36]. By contrast, CA1 ligands such as cholecystokinin (CCK), different types of FGF, and somatostatin (SST) are reported to enhance hippocampal LTP [37–40]. Thyroid hormone deficiency is associated with pronounced deficits in synaptic plasticity (e.g., [41–43]). Caution is urged, however, because some ligands may have distinct (even converse) effects on CA1 versus DG LTP, perhaps pointing to functional differences in the receptors expressed in different hippocampal regions. Nonetheless, based on the published literature, a clear pattern emerges in which challenge ligands (DG) predominantly impair LTP, whereas sufficiency ligands (CA) promote LTP (Figure 5 and Table S8).

**Fig. 5.**
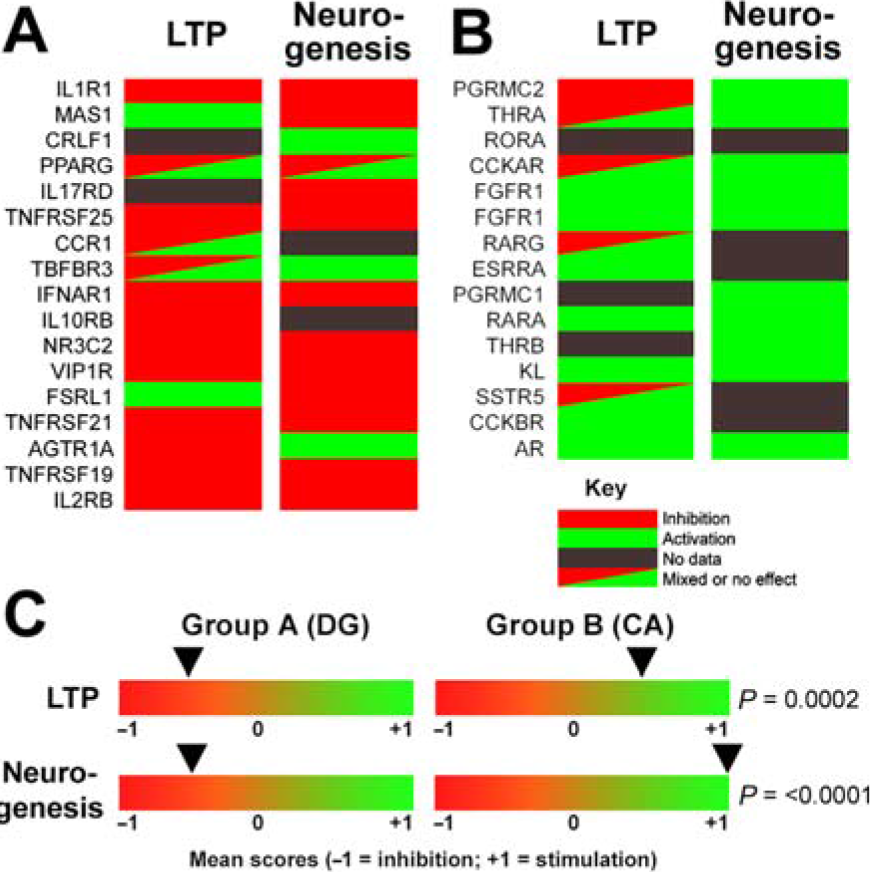
Differential effects of receptor activation on long-term potentiation (LTP) and neurogenesis. (**A**) Group A (DG/challenge). (**B**) Group B (CA/sufficiency). Individual genes are ordered as in Figures 3 and 4. (**C**) Mean scores for the two groups, demonstrating that group A receptors tend to suppress both LTP and neurogenesis, whereas group B receptors tend to promote both parameters. The differential patterns of stimulation/inhibition of LTP and neurogenesis were highly significant between the two groups.

#### Neurogenesis

The literature also records differential effects of DG and CA ligands. Group A (DG/challenge) ligands such as glucocorticoids, interleukins, interferons, and TNF-α are reported to inhibit neurogenesis (e.g., [44–50]) whereas group B (CA/sufficiency) ligands such as estrogen, progesterone, and FGF stimulate neurogenesis (e.g., [51–55]. There are some discordances, particularly when comparing long- and short-term effects (for example for glucocorticoids, reviewed in [47]). However, agents targeting DG predominantly suppress neurogenesis, whereas those targeting CA regions increase neurogenesis (Figure 5 and Table S8).

Because of the small number of samples, differences between each group (DG/CA)/parameter (LTP/neurogenesis) and a random distribution were not uniformly significant (range *P* = 0.005–0.114 for four comparisons and two statistical tests). By contrast, intergroup comparisons revealed that the differences between groups A and B regarding LTP and neurogenesis were consistently highly significant (LTP, *t* test, *P* = 0.0001; chi-square test, *P* = 0.0028; neurogenesis, *t* test, *P* = 0.0002; chi-square test, *P* = 0.006) confirming that the patterns are indeed different.

In conclusion, ligand effects on both LTP and neurogenesis confirm that these hippocampal receptors are functional. Moreover, they indicate that the challenge/sufficiency axis extends to receptor function, wherein DG/challenge receptors predominantly inhibit both neurogenesis and synaptic plasticity, whereas CA/sufficiency ligands principally promote both parameters.

## DISCUSSION

This work confirms and extends prior suggestions that the HPC is involved in internal sensing, as reflected here by greater expression of endocrine receptors than in any other brain region, including CX and CB.

With regard to our first question (how many receptors), we report that 86 of 253 (34%) endocrine receptor genes are expressed in mouse HPC, and 17/98 (17.3%) are exclusively expressed in HPC, values markedly higher than for any other brain region. This accords with our previous data, based on small sample size, that 37% (21–59%, 95% CI) of mouse genes are expressed in HPC, a selection that predominantly encodes endocrine receptors and signaling molecules [21]. Aside from CX and CB, only low-level expression of these receptors was observed in other comparable brain regions (e.g., OLF, thalamus, pons/medulla, pallidum, or striatum; hypothalamus was not studied); these represent ca 4% of all receptors studied.

Thus, of all major brain regions in mouse, endocrine receptor genes are most prominently expressed in HPC, attesting that the present-day HPC is likely to play a sensory role in sensing internal (endocrine) markers of body physiology, arguing that the sensory function attributed to the primeval hippocampus [7–9] has been retained to this day.

Our analysis has focused largely on hormonal ligands, and has not addressed whether the HPC can directly sense levels of low molecular weight ligands (e.g., minerals, pH, CO_2_, etc.) because much less is known about their receptors. For example, NHE4 (SLC9A4), that is activated by hypertonicity, is well expressed in HPC (Allen Brain Atlas), but its exact function is unknown. It could mediate direct sensing of metabolites, although this remains speculative. It is possible that, with evolution, the mouse HPC now responds principally to peripheral hormones that act as proxies for metabolite levels. For example, aldosterone, a salt regulatory hormone, targets NR3C2 in the HPC.

Regarding our second question (patterning within the HPC), we report a highly significant non-random distribution of receptor expression across different HPC subregions of mouse HPC. Receptors whose biological function is known or may be inferred (‘informative’ genes, *n* = 32) were expressed in a highly structured pattern within the formation. Ligands signaling different aspects of challenge (termed here group A: stress, infection, inflammation, blood pressure fall) were principally found to target receptors expressed in DG, whereas ligands signaling aspects of sufficiency (group B: androgens, endocrine FGF, estrogens, progestins, retinoic acid, thyroid hormones) instead principally target the CA regions, with a mean 8.33-fold difference in the DG versus CA expression ratio (*P* < 0.0001).

Although the validity of this distinction remains open to debate (see Results for the underlying rationale), for the purposes of discussion we term this a ‘challenge/sufficiency’ axis. The highly ordered (DG vs CA) segregation of receptor expression in mouse brain raises the question of the function of this segregation (see below).

We also report that the challenge/sufficiency axis accurately mirrors the effects of DG versus CA ligands. With few exceptions, DG/challenge receptors inhibit, whereas CA/sufficiency ligands promote, both neurogenesis and synaptic potentiation.

The contrasting effects on synaptic potentiation suggest that the hippocampus might act as an integrator of positive and negative information: given the paradigmatic hippocampal circuit: cortex → DG → CA3 → CA1 → cortex, the output of the hippocampus is likely to represent the summation of ligand effects on DG and CA regions. The recorded modulation of synaptic potentiation (and thus of overall neurotransmission through the HPC) by endocrine receptor ligands leads us to speculate that the ancestral function of LTP may have been to indicate relevant physiological states worthy of encoding in memory traces, ranging from no LTP (highly adverse context) to potent LTP (highly beneficial context).

A key question concerns whether the challenge/sufficiency axis is reiterated in primates. Preliminary inspection of the microarray-based Allan Human Brain Atlas (http://human.brain-map.org/) fully confirms selective endocrine receptor expression in human HPC, consistent with internal sensing deficits in HPC-ablated patient H.M. [3], but the human data (from elderly individuals) are not strictly comparable to the analyzed data from young mice (and are therefore not presented). There are also hints that DG/CA patterning may be less well conserved in human (not presented), but we note that strict conservation of this patterning across vertebrates is unlikely because, for example, birds and reptiles lack a morphological dentate gyrus (e.g., [56]). Indeed, there is no *a priori* reason why physical segregation of challenge versus sufficiency signaling should be necessary. We suspect that mouse brain may be a special (but informative) case – analysis of this species has pointed, for the first time, to differential HPC receptor localization according to function, providing a new and unexpected perspective on hippocampal function.

Although comprehensive *in situ* receptor expression data in human are so far lacking, there is firm evidence that a functional challenge/sufficiency axis also operates. The human HPC is at the heart of anxiety [57,58], as well as of stress responses and depression. Extensive review would be out of place, but we note that clinical administration of ‘challenge’ ligands (DG in mouse) such as IL-1α, IL-2, IFN-α, IFN-β, and TNF-α produces malaise and sickness behavior [59–64], that has been suggested to be akin to anxiety/depression, whereas ‘sufficiency’ ligands (CA regions in mouse) such as androgens, IGF-1, and thyroid hormone have converse positive effects (e.g., [65–67]), all of which target HPC receptors, indicating that the axis is also functional in human. Systematic inventory of clinical data on challenge/sufficiency ligands will be necessary to confirm this contention.

Nonetheless, we observe an accurate correlation between ligands targeting CA regions and antidepressant/anxiolytic benefits, and the converse for DG ligands. This parallels effects on neurogenesis, where CA ligands predominantly promote neurogenesis in the HPC whereas DG ligands inhibit neurogenesis. This is of special note given that stimulation of HPC neurogenesis has been directly linked to antidepressant action and has been used for new antidepressant drug screening (e.g., [68,69]); differential receptor localization may provide novel indicators for the development of new antidepressants/anxiolytics.

In sum, the selective expression of endocrine receptors in mouse HPC, further highlighted by challenge–sufficiency patterning of endocrine receptor expression, argues that internal sensing remains a core function of the HPC. This accords with evolutionary theory that the HPC arose from a chemosensory epithelium [7–9], and argues that the present-day HPC in particular has retained the ability to monitor the internal milieu of the body. Interoception mediated by the hippocampus may thus provide a new dimension to context-dependent memory encoding, extending from ‘where’ and ‘when’ to ‘how I feel’.

It will be vital to test these concepts in mice genetically engineered to express designer receptors only in DG versus CA regions, and to study the effect of ligand administration on physiology, behavior, and memory. It would also be very informative to study cross-species conservation of expression in larger mammals (rabbit, sheep, non-human primates) where the relative contribution of the hypothalamus (that was too small to be analyzed) could be examined in detail. Moreover, in addition to looking forwards (from mouse to primates), it would be highly illuminating (i) to examine in detail the trajectories of endocrine receptor expression during early development, and (ii) to address the expression profiles of homologs of these genes in other representatives of the vertebrate lineage including birds, reptiles, and fish. One promising line of investigation will be to dissect memory processes in the earliest organisms that encode associations between different internal and external stimuli. Addressing the earliest precedents, and the traces these have left in extant species, will be a fertile territory for new insights into the operation of the human brain.

## Supplementary data

Table S1: List of 250 endocrine receptors studied in this work.

Table S2: Primary endocrine receptor gene expression data.

Table S3: Primary hippocampal endocrine receptor gene expression data.

Table S4: Pairwise correlation analysis.

Table S5: List of informative genes.

Table S6: Comparison of Allen Brain Atlas and HippoSeq expression data.

Table S7: Expression patterns of additional informative receptors confirms the challenge/sufficiency axis.

Table S8: Compilation of ligand effects on LTP and neurogenesis.

Box S1. Observations on the molecular functions of relevant endocrine receptors.

## Acknowledgments

We thank the Allen Brain Institute (Seattle, WA, USA) and Janelia (Ashburn, VA, USA) for making their data publicly available, and to whom we express our deep appreciation. S.S. thanks the Carnegie Trust for the Universities of Scotland for a vacation scholarship. All data needed to evaluate the conclusions in the paper are presented in the paper and/or the supplementary materials online.

## Author Contributions

**Conceptualization:** Richard Lathe, Gernot Riedel.

**Investigation:** Sheena Singadia, Richard Lathe, Gernot Riedel.

**Methodology:** Richard Lathe, Gernot Riedel, Sheena Singadia.

**Statistical analysis:** Crispin Jordan, Richard Lathe

**Project administration:** Gernot Riedel, Richard Lathe.

**Validation:** Richard Lathe; Sheena Singadia, Crispin Jordan, Gernot Riedel.

**Writing – original draft preparation:** Richard Lathe, Gernot Riedel.

**Writing – review & editing:** Richard Lathe; Sheena Singadia, Crispin Jordan, Gernot Riedel.

## Data Availability Statement

All relevant data are within the manuscript and/or supplementary material online.

## Funding

This research received no specific grant from any funding agency in the public, commercial or not-for-profit sectors.

## Competing interests

The authors have declared that no competing interests exist.

