## Supplementary data for "The interoceptive hippocampus: mouse brain endocrine receptor expression highlights a dentate gyrus (DG)–cornu ammonis (CA) challenge– sufficiency axis"

### The primeval internal sensing function of the hippocampus is conserved in mouse

Supplementary data

Table S1: List of 250 endocrine receptors studied in this work.
Table S2: Primary endocrine receptor gene expression data.
Table S3: Primary hippocampal endocrine receptor gene expression data.
Table S4: Pairwise correlation analysis.
Table S5: List of informative genes.
Table S6: Comparison of Allen Brain Atlas and HippoSeq expression data.
Table S7: Expression patterns of additional informative receptors confirms the challenge/sufficiency axis.
Table S8: Compilation of ligand effects on LTP and neurogenesis.
Box S1. Observations on the molecular functions of relevant endocrine receptors.

**Table S1. List of endocrine receptors analyzed**

| | **RECEPTOR** |
|---|---|
| 1 | ACVR1 |
| 2 | ACVR1B |
| 3 | ACVR1C |
| 4 | ACVR2A |
| 5 | ACVR2B |
| 6 | ACVRL1 |
| 7 | ADIPOR1 |
| 8 | ADIPOR2 |
| 9 | ADORA1 |
| 10 | AGTR1A |
| 11 | AHR |
| 12 | AMHR2 |
| 13 | AR |
| 14 | BMPR1B |
| 15 | CALCR |
| 16 | CALCRL |
| 17 | CASR |
| 18 | CCKAR |
| 19 | CCKBR |
| 20 | CCR1 |
| 21 | CCR10 |
| 22 | CCR2 |
| 23 | CCR3 |
| 24 | CCR4 |
| 25 | CCR5 |
| 26 | CCR6 |
| 27 | CCR7 |
| 28 | CCR8 |
| 29 | CCR9 |
| 30 | CD27 |
| 31 | CD40 |
| 32 | CNTFR |
| 33 | CRHR1 |
| 34 | CRHR2 |
| 35 | CRLF1 |
| 36 | CSF1R |
| 37 | CSF2RB |
| 38 | CXCR1 |
| 39 | CXCR2 |
| 40 | CXCR3 |
| 41 | CXCR4 |
| 42 | CXCR5 |

| | |
|---|---|
| 43 | CXCR6 |
| 44 | CXCR7 |
| 45 | CYSTR1 |
| 46 | CYSTR2 |
| 47 | DNER |
| 48 | EBI1 |
| 49 | EDAR2 |
| 50 | EDNRA |
| 51 | EDNRB |
| 52 | EGFEM1 |
| 53 | EGFR |
| 54 | EPOR |
| 55 | ERBB3 |
| 56 | ERBB4 |
| 57 | ESR1 |
| 58 | ESR2 |
| 59 | ESRRA |
| 60 | ESRRB |
| 61 | ESRRG |
| 62 | F2RL1 |
| 63 | FAS |
| 64 | FFAR1 |
| 65 | FFAR2 |
| 66 | FFAR3 |
| 67 | FGFR1 |
| 68 | FGFR2 |
| 69 | FGFR3 |
| 70 | FGFR4 |
| 71 | FZD1 |
| 72 | GALR1 |
| 73 | GALR2 |
| 74 | GALR3 |
| 75 | GCGR |
| 76 | GFRA2 |
| 77 | GHR |
| 78 | GHRHR |
| 79 | GHSR |
| 80 | GIPR |
| 81 | GLP1R |
| 82 | GLP2R |
| 83 | GPER |
| 84 | GPR12 |
| 85 | GRPR1 |
| 86 | HCRTR1 |
| 87 | HCRTR2 |

| | |
|---|---|
| 88 | IFNAR1 |
| 89 | IFNAR2 |
| 90 | IFNGR1 |
| 91 | IFNGR2 |
| 92 | IGF1R |
| 93 | IGF2R |
| 94 | IL10RA |
| 95 | IL10RB |
| 96 | IL11RA1 |
| 97 | IL11RA2 |
| 98 | IL12RB1 |
| 99 | IL12RB2 |
| 100 | IL13RA1 |
| 101 | IL13RA2 |
| 102 | IL15RA |
| 103 | IL17RA |
| 104 | IL17RB |
| 105 | IL17RC |
| 106 | IL17RD |
| 107 | IL18R1 |
| 108 | IL1R1 |
| 109 | IL23R |
| 110 | IL2RA |
| 111 | IL2RB |
| 112 | IL6R |
| 113 | IL7R |
| 114 | INSR |
| 115 | INSRR |
| 116 | KL |
| 117 | KLB |
| 118 | LCT |
| 119 | LDLR |
| 120 | LEPR |
| 121 | LIFR |
| 122 | LNPEP |
| 123 | LPAR6 |
| 124 | LRP1 |
| 125 | LRP1B |
| 126 | LRP2 |
| 127 | LTB4R1 |
| 128 | LTB4R2 |
| 129 | LTBR |
| 130 | MAG |
| 131 | MAS1 |
| 132 | MC1R |

| | |
|---|---|
| 133 | MC2R |
| 134 | MC3R |
| 135 | MC4R |
| 136 | MC5R |
| 137 | MCHR1 |
| 138 | MET |
| 139 | MLNR |
| 140 | MPL |
| 141 | MRGPRA2B |
| 142 | MRGPRB1 |
| 143 | MRGPRB2 |
| 144 | MRGPRB3 |
| 145 | MRGPRB4 |
| 146 | MRGPRB5 |
| 147 | MRGPRB8 |
| 148 | MRGPRD |
| 149 | MRGPRE |
| 150 | MRGPRF |
| 151 | MRGPRG |
| 152 | MRGPRH |
| 153 | MRGPRX1 |
| 154 | MST1R |
| 155 | MTNR1B |
| 156 | NECTIN1 |
| 157 | NECTIN2 |
| 158 | NECTIN3 |
| 159 | NGFR |
| 160 | NMBR |
| 161 | NMU2R |
| 162 | NMUR1 |
| 163 | NMUR2 |
| 164 | NOD1 |
| 165 | NOD2 |
| 166 | NPR1 |
| 167 | NPR2 |
| 168 | NPR3 |
| 169 | NR3C1 |
| 170 | NR3C2 |
| 171 | NR4A3 |
| 172 | NTSR1 |
| 173 | NTSR2 |
| 174 | NURR1 |
| 175 | OGFRL1 |
| 176 | OPRL1 |
| 177 | P2RY1 |

| | |
|---|---|
| 178 | P2RY12 |
| 179 | P2RY2 |
| 180 | PAQR3 |
| 181 | PAQR4 |
| 182 | PAQR5 |
| 183 | PAQR6 |
| 184 | PAQR7 |
| 185 | PAQR8 |
| 186 | PAQR9 |
| 187 | PDGFRA |
| 188 | PDGFRB |
| 189 | PGR |
| 190 | PGRMC1 |
| 191 | PGRMC2 |
| 192 | PILRA |
| 193 | PILRB |
| 194 | PLAUR |
| 195 | PPARA |
| 196 | PPARD |
| 197 | PPARG |
| 198 | PPYR1 |
| 199 | PRLR |
| 200 | PTAFR |
| 201 | PTGIR |
| 202 | PTH1R |
| 203 | PTHR2 |
| 204 | PVR |
| 205 | RARA |
| 206 | RARB |
| 207 | RARG |
| 208 | RELL1 |
| 209 | RELL2 |
| 210 | RELT |
| 211 | RORA |
| 212 | RORB |
| 213 | RXFP1 |
| 214 | RXFP2 |
| 215 | RXFP3 |
| 216 | RXFP4 |
| 217 | SCTR |
| 218 | SLC9A4 |
| 219 | SORL1 |
| 220 | SSTR1 |
| 221 | SSTR2 |
| 222 | SSTR4 |

| | |
|---|---|
| 223 | SSTR5 |
| 224 | TACR1 |
| 225 | TACR2 |
| 226 | TGFBR1 |
| 227 | TGFBR2 |
| 228 | TGFBR3 |
| 229 | THRA |
| 230 | THRB |
| 231 | TNFRSF10B |
| 232 | TNFRSF11A |
| 233 | TNFRSF11B |
| 234 | TNFRSF12A |
| 235 | TNFRSF13C |
| 236 | TNFRSF14 |
| 237 | TNFRSF19 |
| 238 | TNFRSF1A |
| 239 | TNFRSF21 |
| 240 | TNFRSF22 |
| 241 | TNFRSF23 |
| 242 | TNFRSF25 |
| 243 | TNFRSF26 |
| 244 | TNFRSF8 |
| 245 | TNFRSF9 |
| 246 | TRHR |
| 247 | TSHR |
| 248 | TSHRB |
| 249 | TTRHR2 |
| 250 | VDR |
| 251 | VIPR1 |
| 252 | VIPR2 |
| 253 | VMN1R234 |

**Table S2. Primary endocrine receptor expression data**
(ImageJ scanning arbitrary units)

| RECEPTOR | HPC | CB | CX |
|---|---|---|---|
| ACVR1 | 30.51 | 0.00 | 2.31 |
| ACVR1B | 12.75 | 2.87 | 11.34 |
| ACVR1C | 0.00 | 0.00 | 0.00 |
| ACVR2A | 7.22 | 0.00 | 0.00 |
| ACVR2B | 0.00 | 0.00 | 0.00 |
| ACVRL1 | 0.00 | 0.00 | 7.69 |
| ADIPOR1 | 6.97 | 3.17 | 2.77 |
| ADIPOR2 | 0.00 | 5.31 | 0.00 |
| ADORA1 | 13.72 | 0.00 | 0.00 |
| AGTR1A | 16.20 | 15.91 | 31.26 |
| AHR | 0.00 | 0.00 | 0.00 |
| AMHR2 | 0.00 | 0.00 | 0.00 |
| AR | 2.69 | 0.00 | 0.00 |
| BMPR1B | 0.00 | 0.00 | 0.00 |
| CALCR | 0.00 | 0.00 | 0.00 |
| CALCRL | 0.00 | 0.00 | 0.00 |
| CASR | 0.00 | 0.00 | 0.00 |
| CCKAR | 19.52 | 7.77 | 8.60 |
| CCKBR | 6.14 | 0.00 | 13.16 |
| CCR1 | 27.87 | 14.40 | 29.67 |
| CCR10 | 0.00 | 0.00 | 0.00 |
| CCR2 | 0.00 | 0.00 | 0.00 |
| CCR3 | 0.00 | 0.00 | 0.00 |
| CCR4 | 0.00 | 0.00 | 0.00 |
| CCR5 | 0.00 | 0.00 | 0.00 |
| CCR6 | 0.00 | 0.00 | 0.00 |
| CCR7 | 0.00 | 0.00 | 0.00 |
| CCR8 | 0.00 | 0.00 | 0.00 |
| CCR9 | 0.00 | 0.00 | 0.00 |
| CD27 | 0.00 | 0.00 | 0.00 |
| CD40 | 13.13 | 0.00 | 0.00 |
| CNTFR | 16.52 | 24.71 | 29.26 |
| CRHR1 | 0.00 | 15.21 | 7.70 |
| CRHR2 | 0.00 | 0.00 | 0.00 |
| CRLF1 | 36.14 | 0.00 | 0.00 |
| CSF1R | 16.65 | 10.09 | 10.43 |
| CSF2RB | 0.00 | 0.00 | 0.00 |
| CXCR1 | 0.00 | 0.00 | 0.00 |
| CXCR2 | 0.00 | 0.00 | 0.00 |
| CXCR3 | 0.00 | 0.00 | 0.00 |
| CXCR4 | 0.00 | 0.00 | 0.00 |
| CXCR5 | 0.00 | 0.00 | 0.00 |
| CXCR6 | 0.00 | 0.00 | 0.00 |

| | | | |
|---|---|---|---|
| CXCR7 | 0.00 | 0.00 | 0.00 |
| CYSTR1 | 0.00 | 0.00 | 0.00 |
| CYSTR2 | 0.00 | 0.00 | 0.00 |
| DNER | 24.06 | 20.25 | 7.94 |
| EBI1 | 0.00 | 0.00 | 0.00 |
| EDAR2 | 0.00 | 0.00 | 0.00 |
| EDNRA | 0.00 | 0.00 | 0.00 |
| EDNRB | 0.00 | 12.70 | 0.00 |
| EGFEM1 | 18.43 | 0.00 | 0.00 |
| EGFR | 0.00 | 0.00 | 0.00 |
| EPOR | 18.43 | 0.00 | 13.65 |
| ERBB3 | 0.00 | 0.00 | 0.00 |
| ERBB4 | 8.52 | 0.00 | 10.19 |
| ESR1 | 0.00 | 0.00 | 0.00 |
| ESR2 | 0.00 | 0.00 | 0.00 |
| ESRRA | 10.59 | 11.30 | 2.11 |
| ESRRB | 0.00 | 0.00 | 0.00 |
| ESRRG | 0.00 | 0.00 | 0.00 |
| F2RL1 | 13.42 | 0.00 | 11.51 |
| FAS | 0.00 | 0.00 | 0.00 |
| FFAR1 | 0.00 | 0.00 | 0.00 |
| FFAR2 | 0.00 | 0.00 | 0.00 |
| FFAR3 | 0.00 | 0.00 | 0.00 |
| FGFR1 | 22.33 | 9.85 | 0.00 |
| FGFR2 | 0.00 | 9.73 | 6.47 |
| FGFR3 | 2.29 | 5.90 | 2.94 |
| FGFR4 | 0.00 | 0.00 | 0.00 |
| FZD1 | 11.84 | 9.96 | 1.53 |
| GALR1 | 0.00 | 0.00 | 0.00 |
| GALR2 | 0.00 | 0.00 | 0.00 |
| GALR3 | 2.61 | 5.50 | 9.72 |
| GCGR | 0.00 | 0.00 | 0.00 |
| GFRA2 | 1.03 | 7.23 | 24.50 |
| GHR | 0.00 | 0.00 | 0.00 |
| GHRHR | 0.00 | 0.00 | 0.00 |
| GHSR | 0.00 | 0.00 | 0.00 |
| GIPR | 0.00 | 0.00 | 0.00 |
| GLP1R | 0.00 | 0.00 | 0.00 |
| GLP2R | 0.00 | 0.00 | 11.56 |
| GPER | 0.00 | 0.00 | 0.00 |
| GPR12 | 10.88 | 0.00 | 0.00 |
| GRPR1 | 0.00 | 0.00 | 0.00 |
| HCRTR1 | 6.42 | 0.00 | 0.00 |
| HCRTR2 | 0.00 | 0.00 | 0.00 |
| IFNAR1 | 26.14 | 26.54 | 35.09 |
| IFNAR2 | 0.00 | 0.00 | 0.00 |
| IFNGR1 | 0.00 | 0.00 | 4.28 |
| IFNGR2 | 13.97 | 0.00 | 0.00 |

| | | | |
|---|---|---|---|
| IGF1R | 21.03 | 9.04 | 1.57 |
| IGF2R | 7.33 | 5.45 | 10.16 |
| IL10RA | 0.00 | 0.00 | 0.00 |
| IL10RB | 47.48 | 13.47 | 29.79 |
| IL11RA1 | 6.33 | 1.61 | 3.20 |
| IL11RA2 | 0.00 | 0.00 | 8.66 |
| IL12RB1 | 0.00 | 0.00 | 0.00 |
| IL12RB2 | 0.00 | 0.00 | 0.00 |
| IL13RA1 | 0.00 | 0.00 | 0.00 |
| IL13RA2 | 0.00 | 0.00 | 0.00 |
| IL15RA | 0.00 | 0.00 | 0.00 |
| IL17RA | 0.00 | 0.00 | 11.34 |
| IL17RB | 0.00 | 0.00 | 0.00 |
| IL17RC | 0.00 | 0.00 | 0.00 |
| IL17RD | 32.79 | 0.10 | 9.18 |
| IL18R1 | 0.00 | 0.00 | 0.00 |
| IL1R1 | 11.60 | 0.00 | 0.00 |
| IL23R | 0.00 | 0.00 | 0.00 |
| IL2RA | 0.00 | 0.00 | 0.00 |
| IL2RB | 12.17 | 2.47 | 0.10 |
| IL6R | 0.00 | 0.00 | 0.00 |
| IL7R | 0.00 | 0.00 | 0.00 |
| INSR | 0.00 | 0.00 | 0.00 |
| INSRR | 0.00 | 0.00 | 0.00 |
| KL | 28.77 | 7.80 | 5.11 |
| KLB | 0.00 | 0.00 | 0.00 |
| LCT | 23.65 | 0.00 | 0.00 |
| LDLR | 0.00 | 0.00 | 0.00 |
| LEPR | 0.00 | 0.00 | 0.00 |
| LIFR | 10.80 | 3.31 | 11.91 |
| LNPEP | 39.89 | 48.81 | 26.99 |
| LPAR6 | 0.00 | 0.00 | 0.00 |
| LRP1 | 34.69 | 8.75 | 23.84 |
| LRP1B | 8.87 | 0.00 | 1.01 |
| LRP2 | 0.00 | 0.00 | 0.00 |
| LTB4R1 | 0.00 | 0.00 | 0.00 |
| LTB4R2 | 0.00 | 0.00 | 0.00 |
| LTBR | 17.55 | 8.67 | 7.51 |
| MAG | 8.36 | 17.47 | 17.59 |
| MAS1 | 24.37 | 0.00 | 0.00 |
| MC1R | 0.00 | 0.00 | 0.00 |
| MC2R | 12.03 | 0.00 | 0.00 |
| MC3R | 0.00 | 0.00 | 0.00 |
| MC4R | 0.00 | 0.00 | 0.00 |
| MC5R | 0.00 | 0.00 | 0.00 |
| MCHR1 | 0.00 | 0.00 | 0.00 |
| MET | 0.00 | 0.00 | 0.00 |

| | | | |
|---|---|---|---|
| MLNR | 0.00 | 0.00 | 0.00 |
| MPL | 0.00 | 0.00 | 0.00 |
| MRGPRA2B | 0.00 | 0.00 | 0.00 |
| MRGPRB1 | 0.00 | 0.00 | 0.00 |
| MRGPRB2 | 0.00 | 0.00 | 0.00 |
| MRGPRB3 | 0.00 | 0.00 | 0.00 |
| MRGPRB4 | 0.00 | 0.00 | 0.00 |
| MRGPRB5 | 0.00 | 0.00 | 0.00 |
| MRGPRB8 | 0.00 | 0.00 | 0.00 |
| MRGPRD | 0.00 | 0.00 | 0.00 |
| MRGPRE | 0.00 | 0.00 | 0.00 |
| MRGPRF | 0.00 | 0.00 | 0.00 |
| MRGPRG | 0.00 | 0.00 | 0.00 |
| MRGPRH | 0.00 | 0.00 | 0.00 |
| MRGPRX1 | 0.00 | 0.00 | 0.00 |
| MST1R | 0.00 | 0.00 | 0.00 |
| MTNR1B | 0.00 | 0.00 | 0.00 |
| NECTIN1 | 38.35 | 13.10 | 7.10 |
| NECTIN2 | 17.35 | 8.03 | 9.04 |
| NECTIN3 | 21.95 | 0.00 | 23.77 |
| NGFR | 0.00 | 0.00 | 0.00 |
| NMBR | 0.00 | 0.00 | 0.00 |
| NMU2R | 0.00 | 0.00 | 0.00 |
| NMUR1 | 0.00 | 0.00 | 0.00 |
| NMUR2 | 0.00 | 0.00 | 0.00 |
| NOD1 | 0.00 | 0.00 | 0.00 |
| NOD2 | 0.00 | 0.00 | 0.00 |
| NPR1 | 0.00 | 9.20 | 0.00 |
| NPR2 | 25.20 | 12.88 | 8.88 |
| NPR3 | 5.08 | 0.00 | 2.20 |
| NR3C1 | 12.40 | 0.00 | 3.83 |
| NR3C2 | 11.13 | 0.00 | 0.00 |
| NR4A3 | 17.72 | 0.00 | 0.00 |
| NTSR1 | 0.00 | 0.00 | 0.00 |
| NTSR2 | 0.00 | 0.00 | 0.00 |
| NURR1 | 11.31 | 23.54 | 13.67 |
| OGFRL1 | 13.01 | 7.81 | 1.33 |
| OPRL1 | 10.77 | 0.00 | 6.52 |
| P2RY1 | 0.00 | 0.00 | 0.00 |
| P2RY12 | 2.05 | 0.00 | 3.12 |
| P2RY2 | 4.21 | 0.00 | 0.00 |
| PAQR3 | 0.00 | 0.00 | 0.00 |
| PAQR4 | 12.47 | 6.97 | 6.75 |
| PAQR5 | 0.00 | 0.00 | 0.00 |
| PAQR6 | 0.00 | 6.38 | 0.00 |
| PAQR7 | 22.27 | 11.65 | 28.14 |
| PAQR8 | 14.38 | 10.25 | 1.60 |
| PAQR9 | 24.86 | 0.00 | 1.13 |

| | | | |
|---|---|---|---|
| PDGFRA | 3.50 | 2.82 | 4.99 |
| PDGFRB | 1.50 | 2.24 | 2.27 |
| PGR | 0.00 | 0.00 | 0.00 |
| PGRMC1 | 19.39 | 0.90 | 2.05 |
| PGRMC2 | 25.26 | 9.84 | 12.94 |
| PILRA | 0.00 | 0.00 | 0.00 |
| PILRB | 0.00 | 0.00 | 0.00 |
| PLAUR | 0.00 | 0.00 | 0.00 |
| PPARA | 0.00 | 0.00 | 0.00 |
| PPARD | 0.00 | 0.00 | 0.00 |
| PPARG | 47.40 | 18.18 | 32.09 |
| PPYR1 | 0.00 | 0.00 | 0.00 |
| PRLR | 0.00 | 0.00 | 0.00 |
| PTAFR | 0.00 | 0.00 | 0.00 |
| PTGIR | 11.36 | 25.40 | 2.83 |
| PTH1R | 0.00 | 0.00 | 0.00 |
| PTHR2 | 0.00 | 0.00 | 0.00 |
| PVR | 36.91 | 46.29 | 36.21 |
| RARA | 12.70 | 6.86 | 1.20 |
| RARB | 0.00 | 0.00 | 0.00 |
| RARG | 24.58 | 0.00 | 11.18 |
| RELL1 | 0.00 | 0.00 | 0.00 |
| RELL2 | 0.00 | 0.00 | 0.00 |
| RELT | 0.00 | 0.00 | 0.00 |
| RORA | 2.51 | 21.62 | 19.72 |
| RORB | 0.00 | 0.00 | 24.62 |
| RXFP1 | 0.92 | 0.00 | 4.61 |
| RXFP2 | 0.00 | 0.00 | 0.00 |
| RXFP3 | 3.86 | 0.00 | 0.00 |
| RXFP4 | 0.00 | 0.00 | 0.00 |
| SCTR | 0.00 | 0.00 | 0.00 |
| SLC9A4 | 11.99 | 0.00 | 0.56 |
| SORL1 | 50.64 | 18.79 | 36.91 |
| SSTR1 | 0.00 | 0.00 | 0.00 |
| SSTR2 | 0.00 | 0.00 | 5.47 |
| SSTR4 | 19.61 | 0.00 | 4.64 |
| SSTR5 | 0.00 | 0.00 | 0.00 |
| TACR1 | 0.00 | 0.00 | 0.00 |
| TACR2 | 0.00 | 0.00 | 0.00 |
| TGFBR1 | 0.00 | 0.00 | 0.00 |
| TGFBR2 | 0.00 | 1.42 | 5.42 |
| TGFBR3 | 21.02 | 7.65 | 3.75 |
| THRA | 31.17 | 0.00 | 16.20 |
| THRB | 17.55 | 0.00 | 11.70 |
| TNFRSF10B | 0.00 | 0.00 | 0.00 |
| TNFRSF11A | 0.00 | 0.00 | 0.00 |
| TNFRSF11B | 0.00 | 0.00 | 0.00 |
| TNFRSF12A | 0.00 | 0.00 | 0.00 |

| | | | |
|---|---|---|---|
| TNFRSF13C | 0.00 | 0.00 | 0.00 |
| TNFRSF14 | 0.00 | 0.00 | 0.00 |
| TNFRSF19 | 11.17 | 0.00 | 4.36 |
| TNFRSF1A | 0.00 | 0.00 | 0.00 |
| TNFRSF21 | 0.00 | 0.00 | 0.00 |
| TNFRSF22 | 0.00 | 0.00 | 0.00 |
| TNFRSF23 | 0.00 | 0.00 | 0.00 |
| TNFRSF25 | 19.71 | 0.00 | 3.87 |
| TNFRSF26 | 0.00 | 0.00 | 0.00 |
| TNFRSF8 | 0.00 | 0.00 | 0.00 |
| TNFRSF9 | 0.00 | 0.00 | 0.00 |
| TRHR | 0.00 | 0.00 | 0.00 |
| TSHR | 0.00 | 0.00 | 0.00 |
| TSHRB | 0.00 | 0.00 | 0.00 |
| TTRHR2 | 0.00 | 0.00 | 0.00 |
| VDR | 0.00 | 0.00 | 0.00 |
| VIPR1 | 11.21 | 0.00 | 8.86 |
| VIPR2 | 0.00 | 0.00 | 0.00 |
| VMN1R234 | 23.15 | 0.00 | 0.00 |

**Table S3. Primary hippocampal subregion informative receptor expression data**

| Receptor | CA1 | CA2 | CA3 | DG | CA1 | CA2 | CA3 | DG | Group |
|---|---|---|---|---|---|---|---|---|---|
| IL1R1 | 0 | 0 | 0 | 1 | 0 | 0 | 0 | 100 | A |
| MAS1 | 0.037 | 0.44 | 0.464 | 1 | 3.7 | 44 | 46.4 | 100 | |
| CRLF1 | 0.265 | 0.052 | 0.392 | 1 | 26.5 | 5.2 | 39.2 | 100 | |
| PPARG | 0.282 | 0.367 | 0.417 | 1 | 28.2 | 36.7 | 41.7 | 100 | |
| IL17RD | 0.3 | 0.2 | 0.2 | 1 | 30 | 20 | 20 | 100 | |
| TNFRSF25 | 0.361 | 0.322 | 0.286 | 1 | 36.1 | 32.2 | 28.6 | 100 | |
| CCR1 | 0.526 | 0.505 | 0.57 | 1 | 52.6 | 50.5 | 57 | 100 | |
| TGFBR3 | 0.565 | 0.328 | 0.442 | 1 | 56.5 | 32.8 | 44.2 | 100 | |
| IFNAR1 | 0.6 | 0.6 | 0.6 | 1 | 60 | 60 | 60 | 100 | |
| IL10RB | 0.788 | 0.71 | 0.518 | 1 | 78.8 | 71 | 51.8 | 100 | |
| NR3C2 | 0.685 | 0.763 | 0.671 | 1 | 68.5 | 76.3 | 67.1 | 100 | |
| VIPR1 | 0.626 | 0.183 | 0.896 | 1 | 62.6 | 18.3 | 89.6 | 100 | |
| F2RL1 | 0.589 | 1 | 0.067 | 0.879 | 58.9 | 100 | 6.7 | 87.9 | |
| TNFRSF21 | 0.767 | 0.849 | 0.893 | 1 | 76.7 | 84.9 | 89.3 | 100 | |
| AGTR1A | 0.884 | 0.836 | 0.77 | 1 | 88.4 | 83.6 | 77 | 100 | |
| TNFRSF19 | 1 | 0.509 | 0.97 | 0.846 | 100 | 50.9 | 97 | 84.6 | |
| IL2RB | 0.932 | 0.168 | 1 | 0.77 | 93.2 | 16.8 | 100 | 77 | |
| PGRMC2 | 0.377 | 0.323 | 1 | 0.276 | 37.7 | 32.3 | 100 | 27.6 | B |
| THRA | 0.432 | 0.658 | 1 | 0.244 | 43.2 | 65.8 | 100 | 24.4 | |
| RORA | 0.574 | 1 | 0.915 | 0.246 | 57.4 | 100 | 91.5 | 24.6 | |
| CCKAR | 0.8 | 1 | 0.91 | 0.75 | 80 | 100 | 91 | 75 | |
| FGFR1 | 0.78 | 1 | 0.884 | 0.53 | 78 | 100 | 88.4 | 53 | |
| FGFR3 | 0.1 | 1 | 0.467 | 0.033 | 10 | 100 | 46.7 | 3.3 | |
| RARG | 1 | 0.697 | 0.601 | 0.599 | 100 | 69.7 | 60.1 | 59.9 | |
| ESRRA | 1 | 0.309 | 0.58 | 0.638 | 100 | 30.9 | 58 | 63.8 | |
| PGRMC1 | 1 | 0.485 | 0.749 | 0.592 | 100 | 48.5 | 74.9 | 59.2 | |

| | | | | | | | | |
|---|---|---|---|---|---|---|---|---|
| RARA | 1 | 0.552 | 0.54 | 0.311 | 100 | 55.2 | 54 | 31.1 |
| THRB | 1 | 0.526 | 0.237 | 0.203 | 100 | 52.6 | 23.7 | 20.3 |
| KL | 1 | 0.039 | 0.221 | 0.2 | 100 | 3.9 | 22.1 | 20 |
| SSTR4 | 1 | 0.293 | 0 | 0 | 100 | 29.3 | 0 | 0 |
| CCKBR | 1 | 0 | 0 | 0 | 100 | 0 | 0 | 0 |
| AR | 1 | 0 | 0 | 0 | 100 | 0 | 0 | 0 |

**Table S4. Cross-Correlation Analysis**

95% CI values for correlation of normalized (and arcsine square root transformed) gene expression between regions CA1, CA2, CA3, and DG, showing significant positive correlations between CA2 and CA3, and significant negative correlations between CA1 and DG (Bold Font).[a]

| | CA2 | CA3 | DG |
|---|---|---|---|
| CA1 | −0.1652, 0.3194 | −0.2902, 0.2440 | **−0.5067, −0.0990** |
| CA2 | | **0.1991, 0.6087** | −0.1730, 0.2813 |
| CA3 | | | −0.3727, 0.1771 |

[a]In most comparisons, non-normalized gene expression did not differ significantly among regions CA1, CA2, CA3 and DG (paired *t* tests (df = 85): CA1 and CA2: $t = 2.5221$, $P = 0.01353$; CA1 and CA3: $t = 1.2136$, $P = 0.2283$; CA1 and DG: $t = -1.6319$, $P = 0.1064$; CA2 and CA3: $t = -1.2556$, $P = 0.2127$; CA2 and DG: $t = -3.3113$, $P = 0.001364$; CA3 and DG: $t = -2.4651$, $P = 0.01571$); only the difference between CA2 and DG remained significant after accounting for multiple comparisons, with higher gene expression in DG.

**Table S5. List of 'Informative' Endocrine Receptors**[a]

| Receptor | Full name | Known ligands |
|---|---|---|
| IL1R1 | Interleukin 1 receptor type 1 | IL-1α, IL-1β, IL-1RN |
| MAS1 | MAS1 proto-oncogene GPCR | Angiotensin (1–7) |
| CRLF1 | Cytokine receptor-like factor 1 | Cardiotrophin-like cytokine (CLC); the CRLF1/CLC complex activates the ciliary neurotrophic factor receptor (CNTFR) |
| PPARG | Peroxisome proliferator-activated receptor γ | Prostaglandin J2 (15d-PGJ2); 5-oxo-15(S)-HETE and 5-oxo-ETE |
| IL17RD | Interleukin 17 receptor D | IL-17 |
| TNFRSF25 | TNF receptor superfamily 25 | TNF-α/TNFSF12 |
| CCR1 | C-C motif chemokine receptor 1 | Macrophage inflammatory protein 1α (MIP-1α), RANTES, MCP-3, and MPIF-1. |
| TGFBR3 | Transforming growth factor β receptor 3 | TGF-β superfamily members |
| IFNAR1 | Inferferon α/β receptor subunit 1 | Type I interferons, including IFNA, IFNB1, and IFNW1 |
| IL10RB | Interleukin 10 receptor subunit β | IL-10, IL-22, IL-26, IL-28, and IFN-L1 |
| NR3C2 | Nuclear receptor subfamily 3C2; mineralocorticoid receptor (MR) | In the absence of HSD11B expression, corticosterone (mouse)/cortisol (human) is the principal ligand (Box 1) |
| VIPR1 | Vasoactive intestinal polypeptide receptor 1 | VIP |
| F2RL1 | F2R-like trypsin receptor 1/protease-activated receptor PAR2 | Trypsin/thrombin; activated by local proteolysis |
| TNFRSF21 | TNF receptor superfamily 21 | TNF superfamily members |
| AGTR1A | Angiotensin II receptor type 1 | Angiotensin II (Ang 1–8) |
| TNFRSF19 | TNF receptor superfamily 19 | TNF superfamily members |
| IL2RB | Interleukin 2 receptor subunit β | IL-2 |
| PGRMC2 | Progesterone receptor membrane component 2 | Progesterone/possible sigma-2 receptor by analogy to PGRMC1 |
| THRA | Thyroid hormone receptor α | Thyroid hormone (triiodothyronine, T3, and thyroxine, T4) |
| RORA | RAR-related orphan receptor A | Endogenous ligand presumed to be related to retinoic acid; potential ligands include cholesterols and vitamin D derivatives |
| CCKAR | Cholecystokinin A receptor | Non-sulfated members of the cholecystokinin (CCK) family |
| FGFR1 | Fibroblast growth factor receptor 1 | Inferred to respond to endocrine FGFs including FGF19, FGF21, and FGF23 (Box 1) |
| FGFR3 | Fibroblast growth factor receptor 3 | Inferred to respond to endocrine FGFs such as FGF19, FGF21, and FGF23 (Box 1) |
| RARG | Retinoic acid receptor γ | All-*trans* retinoic acid, 9-*cis* retinoic acid |
| ESRRA | Estrogen-related receptor α | Estrogen-related molecules (Box 1), cholesterols, possibly farnesyl pyrophosphate |
| PGRMC1 | Progesterone receptor membrane component 1 | Progesterone; sigma-2 receptor (e.g., haloperidol) |
| RARA | Retinoic acid receptor α | All-*trans* retinoic acid, 9-*cis* retinoic acid |
| THRB | Thyroid hormone receptor β | Thyroid hormone (triiodothyronine, T3, and thyroxine, T4) |
| KL | Klotho | FGF coreceptor; inferred to respond to endocrine FGFs including FGF19, FGF21, and FGF23 (Box 1) |
| SSTR4 | Somatostatin receptor 4 | Somatostatin peptides (SST-14, SST-28) |
| CCKBR | Cholecystokinin B receptor | Gastrin and cholecystokinin (CCK) |
| AR | Androgen receptor; nuclear receptor superfamily 3C4 | Testosterone, dihydrotestosterone, modified estrogens (e.g., androstenediol; Box 1) |

[a]Receptor order is a gradient, with DG-expressed receptors at the top of the table, and CA-expressed receptors at the foot.

**Table S6. Comparison of the Allen Brain Atlas (ABA) and HippoSeq Analyses of Gene Expression in Mouse Hippocampus**

| | ABA | HippoSeq |
|---|---|---|
| Method | *In situ* hybridization with coding-region probes following formalin fixation | Deep sequencing of transgenically selected marked cells followed by computer matching |
| Mouse strain | Mixture<br>129S1/SvlmJ<br>C57BL/6J<br>CAST/EiJ<br>DBA/2J<br>PWD/PhJ<br>SPRET/EiJ<br>WSB/EiJ | C57BL/6 |
| Sex | Mixture M/F | Mixture M/F |
| Age | 4 Weeks | Postnatal days P25–P32 |
| Hippocampus | Gradient; dorsal selected | Dorsal selected |
| Phase of light/dark cycle | 8 am–noon | Mid light cycle |
| Number of repeats for one receptor | Single animals for each probe | 100 cells per animal, three animals per receptor |
| *n* (animals) | *n* = 253 | *n* = 3 for each region (pooled); for 7 regions *n* = 21 |
| Output | *In situ* hybridization intensity, single probe (or small number), amplitude score (arbitrary units via ImageJ) | FPKM sequence reads; single transgenic expression tag |

**Table S7. Expression Patterns of Additional Informative Receptors Confirms the Challenge/Sufficiency Axis**

| **Full name** | ***Gene*** | **Challenge/ sufficiency** | **Hippocampal expression[a]** |
|---|---|---|---|
| Interleukin 6 receptor | *Il6r* | Challenge | DG >> CA regions |
| Growth hormone secretagogue receptor | *Ghsr* | Challenge | DG >> CA regions |
| Opioid growth factor receptor | *Ogfr* | Challenge | DG >> CA regions |
| Irisin receptor | *Itgav* | Challenge | DG >> CA regions |
| Leptin receptor | *Lepr* | Sufficiency | CA1 = CA3 = DG |
| Glucagon-like peptide 1 receptor | *Glp1r* | Sufficiency | CA3/CA1 >> DG |

[a]Data: Allen Brain Atlas (ABA) and/or HippoSeq. For *Il6r* expression was below the limit of detection in ABA but HippoSeq confirmed restricted expression principally in DG.

**Table S8. Literature survey of ligand/receptor effects on long-term potentiation (LTP) and neurogenesis**

| Receptor | Full name | Ligand(s) | Effects of activation on LTP in DG | CA1 | Refs | Effects on neurogenesis in DG | Refs |
|---|---|---|---|---|---|---|---|
| IL1R1 | Interleukin 1 receptor type 1 | IL-1α | ↓ | | (1) | ↓ | (6) |
| | | IL1-β | ↓ | ↓ | (2-5) | | |
| MAS1 | MAS1 proto-oncogene GPCR | Angiotensin 1–7 | | ↑[d] | (7) | ↓ | (8) |
| CRLF1 | Cytokine receptor-like factor 1 | Cardiotrophin-like cytokine (CLC); the CRLF1/CLC complex activates the ciliary neurotrophic factor receptor (CNTFR) | | | N/L | ↑ | (9-11) |
| PPARG | Peroxisome proliferator-activated receptor gamma | Prostaglandin J2 (15d-PGJ2); 5-oxo-15(S)-HETE and 5-oxo-ETE | | NE | (12) | ↑ (ld) ↓ (hd) | (13) |
| IL17RD | Interleukin 17 receptor D | IL-17 | | | N/L | ↓ | (14) |
| TNFRSF25 | TNF receptor superfamily 25 | TNFSF12 (TNFα) | ↓ | NE | (15,16) (17) | ↓ | (18) |

| Receptor | Full name | Ligand(s) | DG | CA1 | Refs | Effects on neurogenesis in DG | Refs |
|---|---|---|---|---|---|---|---|
| CCR1 | C-C motif chemokine receptor 1 | Macrophage inflammatory protein 1α (MIP-1α – also known as CCL3) | | ↓ | (19) | N/L | |
| | | RANTES | | NE | (20) | | |
| | | MCP-3 and MPIF-1 | | | N/L | | |
| TGFBR3 | Transforming growth factor beta receptor 3 | TGF-β superfamily members | ↓ | ↑ | (21) (22) | ↑ | (23;24) |
| IFNAR1 | Inferferon α/β receptor subunit 1 | Type I interferons, including IFNA, IFNB1, and IFNW1 | | ↓ | (25) | ↓ | (26) |
| IL10RB | Interleukin 10 receptor subunit β | IL10, IL22, IL26, IL28, and IFNL1 | ↓ | | (27) | | N/L |
| NR3C2 | Nuclear receptor subfamily 3C2; mineralocorticoid receptor | In the absence of HSD11B expression, corticosterone (mouse)/cortisol (human) is the principal ligand (see Box 1) | ↓ | ↓ | (28) (29;30) | ↓ | (31;32) |
| VIPR1 | Vasoactive intestinal polypeptide receptor 1 | VIP | | ↓ | (33) | ↓ | (34;35) |
| F2RL1 | F2R-like trypsin receptor 1/protease-activated receptor PAR2 | Trypsin/thrombin; activated by local proteolysis | | ↑ | (36;37;38) | ↓ | (39) |
| TNFRSF21 | TNF receptor superfamily 21 | TNF superfamily members | | | See TNFRSF25 | | N/L |
| AGTR1A | Angiotensin II receptor type 1 | Angiotensin II (Ang 1–8) | ↓ | | (40-45) | ↑ | (46;47) |

| | | | | | | | |
|---|---|---|---|---|---|---|---|
| TNFRSF19 | TNF receptor superfamily 19 | TNF superfamily members | | | See TNFRSF25 | | N/L |
| IL2RB | Interleukin 2 receptor subunit beta | IL-2 | | ↓ | (48) | ↓ | (49) |
| PGRMC2 | Progesterone receptor membrane component 2 | Progesterone/possible sigma-2 receptor by analogy to PGRMC1 | | ↓ | (50) | ↑ | (51;52) |
| THRA | Thyroid hormone receptor alpha | Thyroid hormone (triiodothyronine, T3, and thyroxine, T4) | ↓ | ↑ | (53-55)<br>(56) | ↑ | (57) |
| RORA | RAR related orphan receptor A | Endogenous ligand presumed to be related to retinoic acid; potential ligands include cholesterols and vitamin D derivatives | | | N/L | | N/L |
| CCKAR | Cholecystokinin A receptor | Non-sulfated members of the cholecystokinin (CCK) family | | ↑<br>NE | (58)<br>(59) | ↑ | (60;61) |
| FGFR1 | Fibroblast growth factor receptor 1 | Inferred to respond to endocrine FGFs including FGF19, FGF21, and FGF23 (Box 1) | ↑ | | (62) | ↑ | (62;63) |
| FGFR3 | Fibroblast growth factor receptor 3 | Inferred to respond to endocrine FGFs such as FGF19, FGF21, and FGF23 (Box 1) | | | N/L (for FGF, see above) | ↑ | (63) |
| RARG | Retinoic acid receptor gamma | All-*trans* retinoic acid, 9-*cis* retinoic acid | | NE | (64) | | N/L |

| | | | | | | | |
|---|---|---|---|---|---|---|---|
| ESRRA | Estrogen-related receptor alpha | Estrogen-related molecules (Box 1), cholesterols, possibly farnesyl pyrophosphate | | ↑ | (65;66) | | N/L |
| PGRMC1 | Progesterone receptor membrane component 1 | Progesterone; sigma-2 receptor (e.g., haloperidol) | | | See above for PGRMC2 | ↑ | (67) |
| RARA | Retinoic acid receptor alpha | All-*trans* retinoic acid, 9-*cis* retinoic acid | | ↑ | (68) | ↑ | (69) |
| THRB | Thyroid hormone receptor beta | Thyroid hormone (triiodothyronine, T3, and thyroxine, T4) | | | N/L | ↑ | (70) |
| KL | Klotho | FGF coreceptor; inferred to respond to endocrine FGFs including FGF19, FGF21, and FGF23 (Box 1) | | ↑ | (71) | ↑ | (72) |
| SSTR4 | Somatostatin receptor 4 | Somatostatin peptides (SST-14, SST-28) | ↓[e] | ↑[e] | (73)<br>(74) | | N/L |
| CCKBR | Cholecystokinin B receptor | Gastrin and cholecystokinin (CCK) | | ↑ | (58;59) | | N/L |
| AR | Androgen receptor; nuclear receptor superfamily 3C4 | Testosterone, dihydrotestosterone (androstenediol; Box 1) | ↑ | ↑ | (75)<br>(76;77) | ↑ | (78-80) |

[a]Table summarizes the list of putative agonists for the informative receptors and the effects of ligands that activate the receptors. Data on long-term potentiation as a proxy for synaptic plasticity (other forms were neglected here; also, no discrimination between early and late phase LTP was performed) are mainly derived from work on rat both in vivo and in brain slices; they are extrapolated to murine function as data on mouse LTP are relatively scarce. Only positive data are mentioned, if no arrow is shown, no literature is available.

[b]Receptor order is a gradient as in Figure 3, with DG-expressed receptors at the top of the table, and CA-expressed receptors at the foot.

[c]Key: ↑ indicates enhancement, ↓ indicates reduction/suppression of activity; N/L, no available literature; NE, no effect reported by cited work; ld, low dose; hd, high dose.

[d]See also AGTR1A.

[e]Note difficulty to differentiate between neuronally released somatostatin from interneurons and exogenous somatostatin from blood.
